# Sperm length evolution in relation to body mass is shaped by multiple trade-offs in tetrapods

**DOI:** 10.1101/2023.09.12.557314

**Authors:** L. Koçillari, S. Cattelan, M.B. Rasotto, F. Seno, A. Maritan, A. Pilastro

## Abstract

Sperm size is highly variable across species and is influenced by various factors including fertilization mode, female reproductive traits and sperm competition. Despite considerable efforts, many questions about sperm size variation remain open. Variation in body size may affect sperm size evolution through its influence on these factors, but the extent to which sperm size variation is linked to body mass remains elusive. In this study, we use the general theory of Pareto Optimality to investigate the relationship between sperm size and body mass across tetrapods. We find that tetrapods fall within a triangular-shaped Pareto front in the trait space of body mass and sperm length suggesting that the evolution of sperm size in relation to body size is shaped by trade-offs. We then explore the three main factors predicted to influence sperm size evolution, namely sperm competition, clutch size and genome size. Our results demonstrate that body mass optimally shapes sperm size evolution in tetrapods mainly through its association with sperm competition and clutch size. Finally, we show that the triangular-shaped Pareto front is maintained when tested separately within mammals, birds, endothermic species and internal fertilizers, suggesting that similar evolutionary trade-offs characterize the evolution of sperm size in relation to body size within taxonomic/phylogenetic and functional subgroups of tetrapods. This study provides insights into the evolutionary mechanisms driving interspecific sperm size variation and highlights the importance of considering multiple trade-offs in optimizing reproductive traits.

## Introduction

*“[…] How does sperm form relate to function? How do different fertilisation environments shape sperm phenotypes, such as fresh-versus salt-water, or hot-versus cold blooded reproductive tracts? How does sperm morphology co-vary with body size, or genome size?”* Matt Gage – Nature Ecology and Evolution 5, pages 1064–1065 (2021)

In a recent commentary on the most comprehensive analysis of sperm size variation across the animal kingdom published to date^1^, Matt Gage^2^ concluded that despite intensive research shedding light on the evolution of the large variability in sperm size and form, many questions remain open. Sperm size, and in particular sperm length, the most common measure used for sperm size (to which we will refer hereafter), varies enormously between taxa^1^, ranging from less than 10 μm to nearly 10^5^ μm (in some fruit flies)^3^. The most extensively studied taxa are arthropods and vertebrates, particularly tetrapods. The smallest and longest sperm recorded so far are both found in arthropods, while sperm size in vertebrates does not show the same level of variation, ranging from slightly more than 10 μm (in external fertilizers such as bony fishes and frogs) to approximately 10^3^ μm (in some urodeles)^4^. It has been recently demonstrated on a large taxonomical scale that the fertilization mode represents an important source of variation, with external fertilizers having, on average, shorter sperm than internal fertilizers^1,5^. Moreover, sperm competition (i.e., when sperm from different males compete to fertilize the same eggs^6–8^) leads to the evolution of i) larger testes, and ii) longer sperm^9^ (with exceptions limited to a few taxa reviewed in^10^. The evolution of larger testes in response to sperm competition has received such a broad empirical support that testes mass is commonly used as a proxy for sperm competition^9^. In contrast, positive selection for longer sperm has been just recently recognized as a general trend^9^, probably because the mechanisms leading to the evolution of longer sperm differ between taxa. In small animals, sperm actively interact in the female reproductive tract and larger sperm have a direct competitive advantage by displacing rival male sperm or saturating female storage organs^6,11^. On the other side, in large animals, sperm have to move via a longer female reproductive tract^12^, which favours the evolution of longer sperm via selection for faster sperm swimming speed^13^. As a result, interspecific variation in body size seems crucial in mediating different processes for the evolution of sperm size under equivalent levels of sperm competition^14^.

Variation in body size may also affect sperm size evolution through its influence on other factors. First, body size negatively correlates with mass-specific metabolic rate in animals (hereafter MSMR)^15,16^. MSMR, in turn, is predicted to influence the evolution of sperm length, as lower metabolic rate may limit the production of long sperm due to low cellular activity^17,18^. This hypothesis may explain why small mammals, which have high MSMR, evolve long sperm in response to sperm competition while the same relationship is not observed in large-bodied mammals, which in contrast have lower mass-specific metabolic rate^17,19^. Second, clutch/litter size which covaries with body size within most vertebrate classes (albeit in different directions^20–22^), may influence the ratio of sperm number/size depending on the taxonomic group. In internal fertilizers clutch size is expected to be positively correlated with sperm size e.g.^11,23^, while in external fertilizers, large clutch sizes are expected to favour the production of numerous sperm^24,25^, at the expense of sperm size. Third, body size is positively correlated with genome size, at least in some groups e.g.^26^, which in turn covaries with nuclear and cellular size^27,28^, but see^29^. It is likely that sperm with a bigger genome require a longer flagellum to compensate for the drag caused by the larger sperm head. Finally, variation in body size frequently reflects variation in ecological characteristics related to reproductive strategies and hence levels of sperm competition^30–33^. Overall, several different interacting evolutionary forces are likely to influence the association between body size and sperm size in animals.

Despite the recent increasing interest in the evolution of sperm size, how and to which extent sperm size variation is linked to body mass remains unknown^2^. Here, we hypothesize that the relationship between sperm length and body mass in tetrapods is a result of trade-offs among multiple factors with potential contrasting effects. We tested our hypothesis within the framework of Pareto Optimality Theory, which predicts that in the presence of evolutionary trade-offs across multiple tasks, the phenotypes fall within low-dimensional geometrical patterns, or Pareto fronts, in the trait space^34^. The geometric form of these Pareto fronts depends on the number of tasks in trade-off (see Fig. 1). In the case of two tasks, the Pareto front forms a line segment; for three tasks, it forms a triangle; and for more tasks, the Pareto front is a convex polygon, whose number of vertices correspond to the number of tasks being in trade-off. In the absence of trade-offs, phenotypes are uniformly distributed as clouds of points in the trait space.

**Figure 1.**
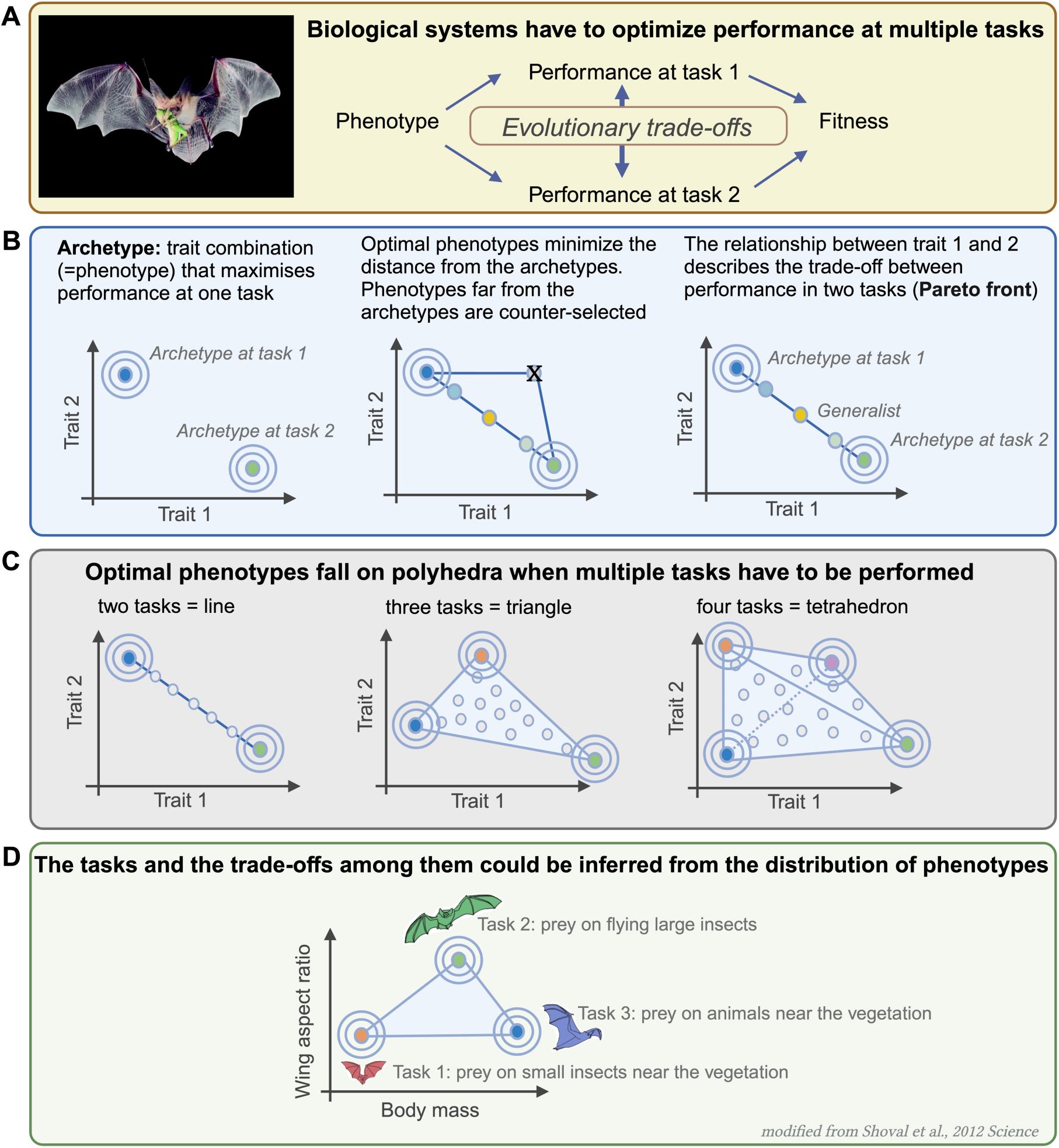
The Pareto Optimality framework. **A)** Biological systems perform multiple tasks to maximize their fitness function. Often, these tasks are in trade-off with each other, making it impossible for a species to optimize performance levels for all tasks simultaneously. The competition between tasks influences the selection of traits, leading organisms to strike a precise balance between traits to maximize their fitness function. **B)** Pareto Optimality is a general framework that predicts that tasks in trade-off lead to low-dimensional distributions of phenotypes, known as Pareto fronts, in the trait space. The vertices of the Pareto fronts are home to archetypal species, or specialists, which perform best at one specific task. The internal part of the Pareto fronts is occupied by generalists, species that are optimal at performing multiple tasks simultaneously, but not as well as the archetypes for individual tasks. **C)** In the two-dimensional trait space, optimal phenotypes fall into convex polygons. For two tasks in trade-off, phenotypes fall into lines; for three tasks, they fall into triangles; for four tasks, they fall into polyhedrons, and so on. **D)** An example of a triangular-shaped Pareto front in the trait space of body mass and wing aspect ratio is shown. The three vertices correspond to archetypes with different predation strategies (edited from^34^)

We investigated multiple, complex trade-offs between tasks using the novel statistical framework of Pareto Task Inference (*ParTI*)^34^. This methodology has been previously been employed in various biological systems, including gene expression^35^, tumours^36^ and proteins^37,38^, trait morphology^34,39–42^, neural systems^43,44^ and behavioural traits^45,46^. We applied ParTI to investigate sperm size variation in relation to body mass in tetrapods, analysing an extensive dataset of 1388 tetrapods, which included 231 amphibians, 115 reptiles, 399 birds, and 643 mammals. We first tested whether the distribution of sperm length and body mass lies on a Pareto front, as expected if multiple trade-offs constrain the evolution of sperm length in association with body mass. We found that tetrapods fall within a triangular-shaped Pareto front. Secondly, we tested the effect of three different factors−clutch/litter size, sperm competition (estimated from testes mass relative to body mass^9^), and genome size−on the shape of sperm length in relation to body mass by conducting a density enrichment analysis on the vertices of the Pareto front. Third, we analysed the sperm length–body mass Pareto front within homogeneous groups based on thermoregulation mode (endothermic versus ectothermic tetrapods), fertilization mode (internal versus external fertilizer species) and at the class level.

## Results

### Tetrapods fall within a triangular-shaped Pareto front in the trait space of body mass and sperm length

To test whether the sperm length-body mass relationship in tetrapods results from trade-offs between multiple tasks, we searched for the presence of Pareto fronts in the space of body mass (BM) and sperm length (SL). We analyzed an extensive dataset of 1388 tetrapods and for each species we collected data on several continuous features. The features include sperm size (µm)^47^, body mass (g), clutch/litter size, relative testes mass as a proxy for sperm competition^9^, and genome size (see Methods). We analysed our dataset using the ParTi algorithm described in^36^ to assess the statistical significance of triangularity for the distribution of data points in the double logarithmic space of body mass and sperm length and found a significant Pareto front (*p* < 0.001, t-ratio test, corrected for False Discovery Rate (FDR) (*p* < 0.005) (Fig. 2) (see Methods).

**Figure 2.**
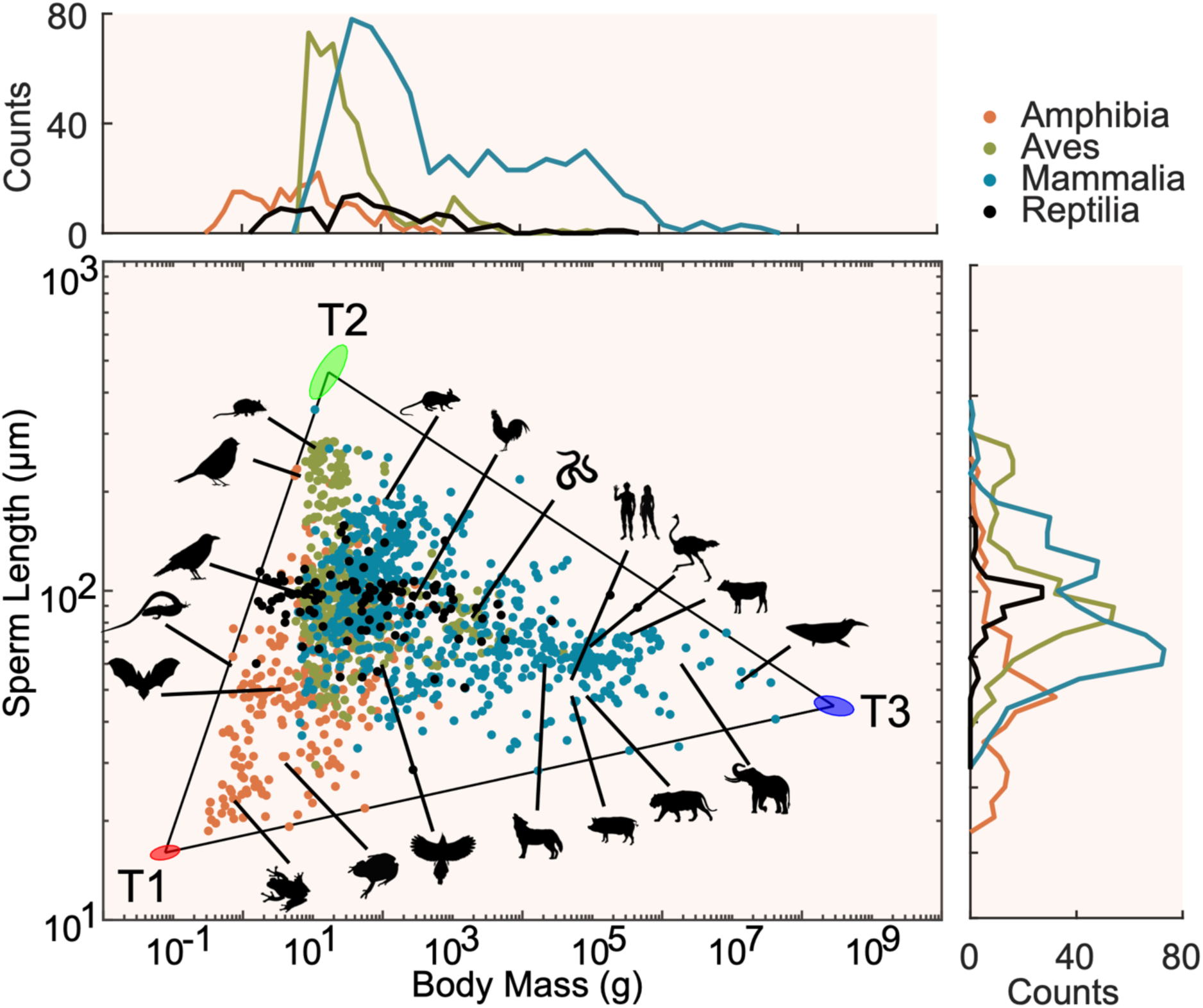
Tetrapods reveal a Pareto front in the log-log space of body mass and sperm length. We pooled together 1388 species from four classes (orange: *Amphibia*, green: *Aves,* blue: *Mammalia,* black: *Reptilia*). Individual species (dots) are plotted in the log-log space of body mass and sperm length. Some species are highlighted by silhouette figures as examples. Species are distributed within a triangular-shaped Pareto front (triangularity-test, *p* = 0.001). The three vertices of the triangle host the archetypal species: vertex T1 (red) is populated by frogs, vertex T2 (green) is populated mainly by birds and small mammals, while vertex T3 (blue) is populated by large mammals. The ellipses represent the error in the vertices positions as determined by bootstrapping. Histograms show the species distributions associated to each class. Silhouette figures were contributed by various authors with a public domain license (public domain mark 1.0; CC0 1.0) from PhyloPic (http://phylopic.org).

We further assessed the goodness of fit of the triangle to the data compared to other convex polygons by computing the sum of squares error as a function of the number of vertices. To do so, we used the Principal Convex Hull/Archetypal analysis (PCHA)^48^ and found that three vertices represent the smallest number that minimize the sum of squares error (Supp. Fig. 1). We finally determined the vertex position using hyperspectral unmixing algorithms, such as Sisal^49^, and obtained the error of the vertex location using bootstrapping^36^ (Supp. Fig. 2).

In conclusion, our results suggest that tetrapods are enclosed within a triangular-shaped Pareto front in the double logarithmic space of body mass and sperm length, indicating that multiple trade-offs have shaped the phenotypic distribution of sperm length in relation to body mass in tetrapods. Furthermore, our results indicate that increasing body size is associated with a moderate increase in sperm length, as indicated by the side of the triangle connecting vertex T1 and T3, which has a slightly positive slope. However, long sperm evolved only in medium-sized tetrapods.

### Species distribution within the Pareto front

We grouped species based on class (Fig. 2) and found that the vertex T1 (red) is exclusively populated by amphibians, which is characterized by species with the lowest body mass and shortest sperm length. In contrast, the vertex T2 (green), which represents species with the longest sperm and small-to-intermediate body mass is populated by species from all classes except reptiles, with a higher frequency of mammals and birds. Mammals with intermediate sperm length and large body mass are located at the vertex T3 (blue). Finally, reptiles are located mainly at the middle of the Pareto front, although they show a relatively large range in both body mass and sperm length. The taxonomic richness of the vertices of the triangle may partly reflect the taxonomic sampling: vertices are less species-dense, and taxonomic groups with smaller sample size are less likely to be represented in the vertices. An alternative, not mutually exclusive explanation, is that factors influencing sperm size evolution in relation to body size are not homogenously distributed within the triangle.

### Clutch size, testes mass and genome size enrich the vertices of the Pareto front

The triangular distribution we found in the trait space of sperm-body size suggests that trade-offs between multiple tasks constrain the evolution of sperm size in relation to body mass. We next explored the properties encoded by the species at the vertices of the triangular distribution. The vertices of the triangle are home of archetypes, i.e., those species that show the highest performance for a specific task (see Methods and Fig. 1). Studying the properties encoded by the species at the vertices is instrumental to inferring the nature of the competing tasks at play. To do so, we considered the three main factors that are predicted by theory (and by the empirical evidence, although at a partial taxonomic level) to influence sperm size evolution: sperm competition, clutch size, and genome size. We used the density enrichment approach to determine the expression of maximal or minimal scores on each feature within the Pareto front.

We examined whether testes mass (as a proxy for the level of sperm competition), clutch size and genome size would enrich near the archetypes (Fig. 3). Our dataset contains clutch size data for 1166 species, testes mass values for 958 species, and genome size data for 331 species (see Table 1). Clutch size is expected to be related to both sperm length and body mass to some extent, and it may be potentially maximized under specific body mass–sperm length combinations (Supp. Fig. 3). Testes mass is expected to be related to body mass and sperm length possibly in contrasting directions (Supp. Fig. 3). Finally, genome size, which is positively associated with cell size, is also expected to influence sperm size variation. To account for the allometric scaling with body mass, we used the standardized residuals for testes mass and clutch size as the main input of the dataset. We found that the density of testes mass and clutch size peaks close to vertex T2 and decreases moving away from it (low testes mass and clutch size for species near vertices T1 and T3, see Fig. 3).

**Figure 3.**
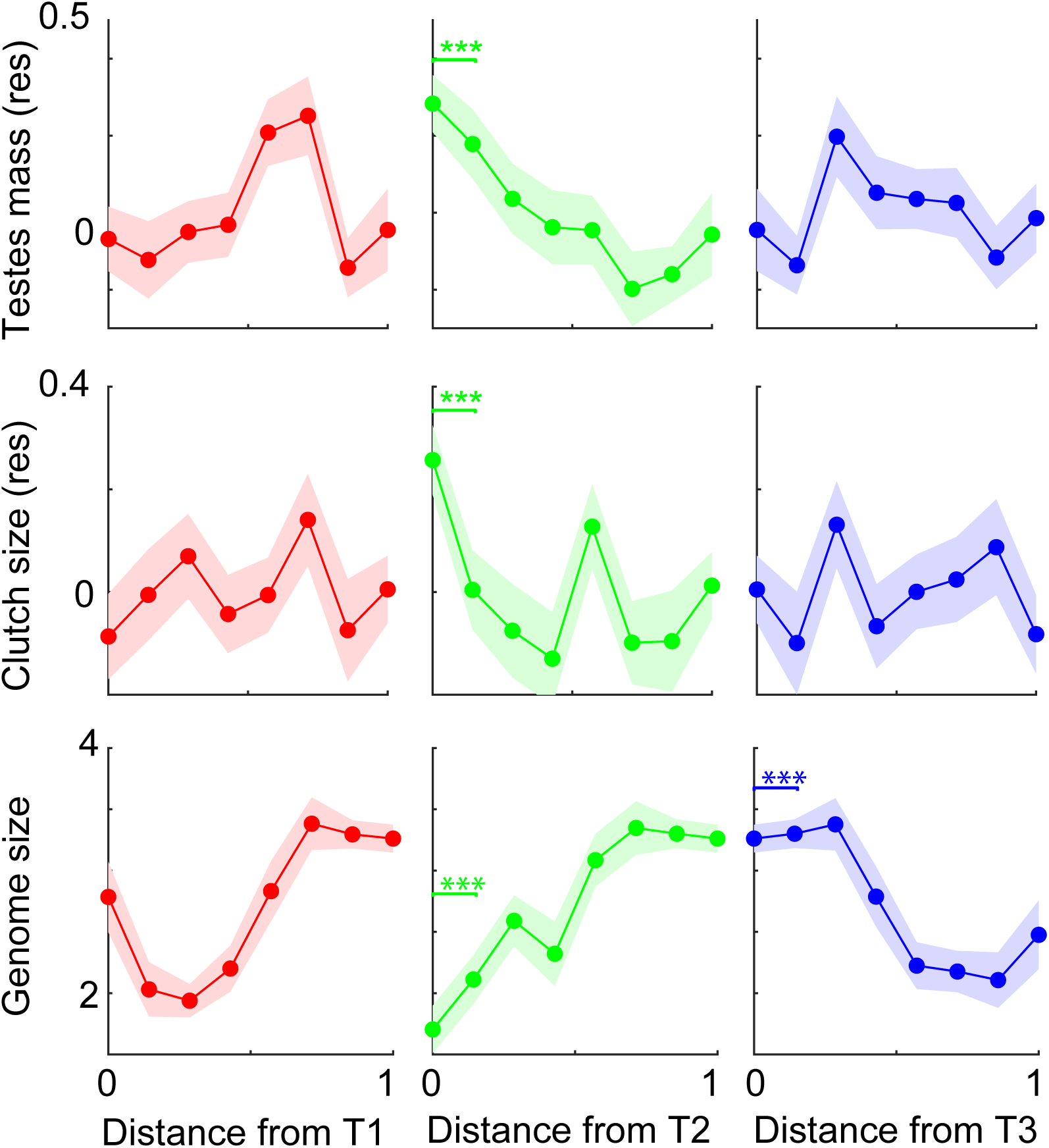
Feature density analysis. We computed the feature density of testes mass (residuals) (top panels), clutch size (residuals) (middle panels), and genome size (bottom panels) as a function of the Euclidean distance from the three vertices of the triangle. Species near vertex T2 show larger clutch size and testes mass, while species close to vertex T3 carry larger genome size. Solid lines represent the mean while shaded areas represent the standard error. Statistical comparisons were performed by running a Wilcoxon rank-sum test (\*\*\**p*<0.001).

**Table 1.**
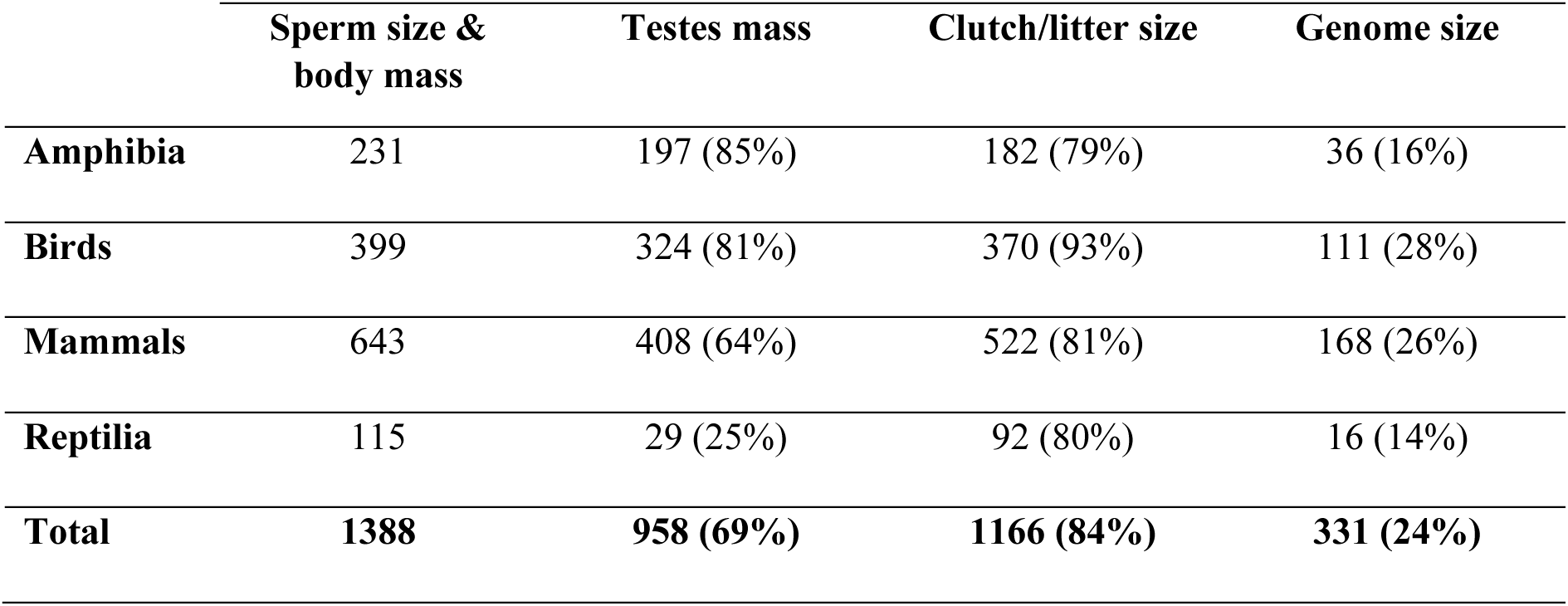
A summary of the number of species with the associated sperm length, body mass, testes mass, clutch/litter size and genome size for each class. In brackets the proportion of species with testes mass, clutch/litter size, and genome size in relation to sperm size and body mass is reported.

These findings reveal that species with long sperm have both large testes mass and clutch size. The size of the genome is instead maximized at the T3 vertex, which corresponds to animals with intermediate sperm sizes and large body mass, and it is minimized at the T2 vertex, which corresponds to species with long sperm sizes and small-to-intermediate body masses.

### Robustness of the Pareto front in relation to thermoregulation and fertilization modes

We next investigated the influence of thermoregulation and fertilization modes on shaping the triangular Pareto front in the space of sperm length and body mass. We thus performed two separate analyses by grouping the species i) based on whether they were endotherms or ectotherms, and ii) based on their fertilization mode, i.e., being an external or internal fertilizer. We found that both endotherms and internal fertilizer species generate significant triangular-shaped Pareto fronts (t-ratio test, *p* = 0.004, *p =* 0.012 respectively), whereas ectotherms and external fertilizer species do not (*p* > 0.05) (Fig.4). We next performed density enrichment analyses on the vertices of the triangles and found that, similar to what we found in the main triangular Pareto front, the clutch size is maximally enriched close to the vertex T2, which is populated by species with long sperm sizes and intermediate body masses (Fig. 5). Our results suggest that the evolutionary constraints shaping sperm size–body mass distribution in tetrapods are robust also within functional groups, namely endotherms, and internal fertilizers.

**Figure 4.**
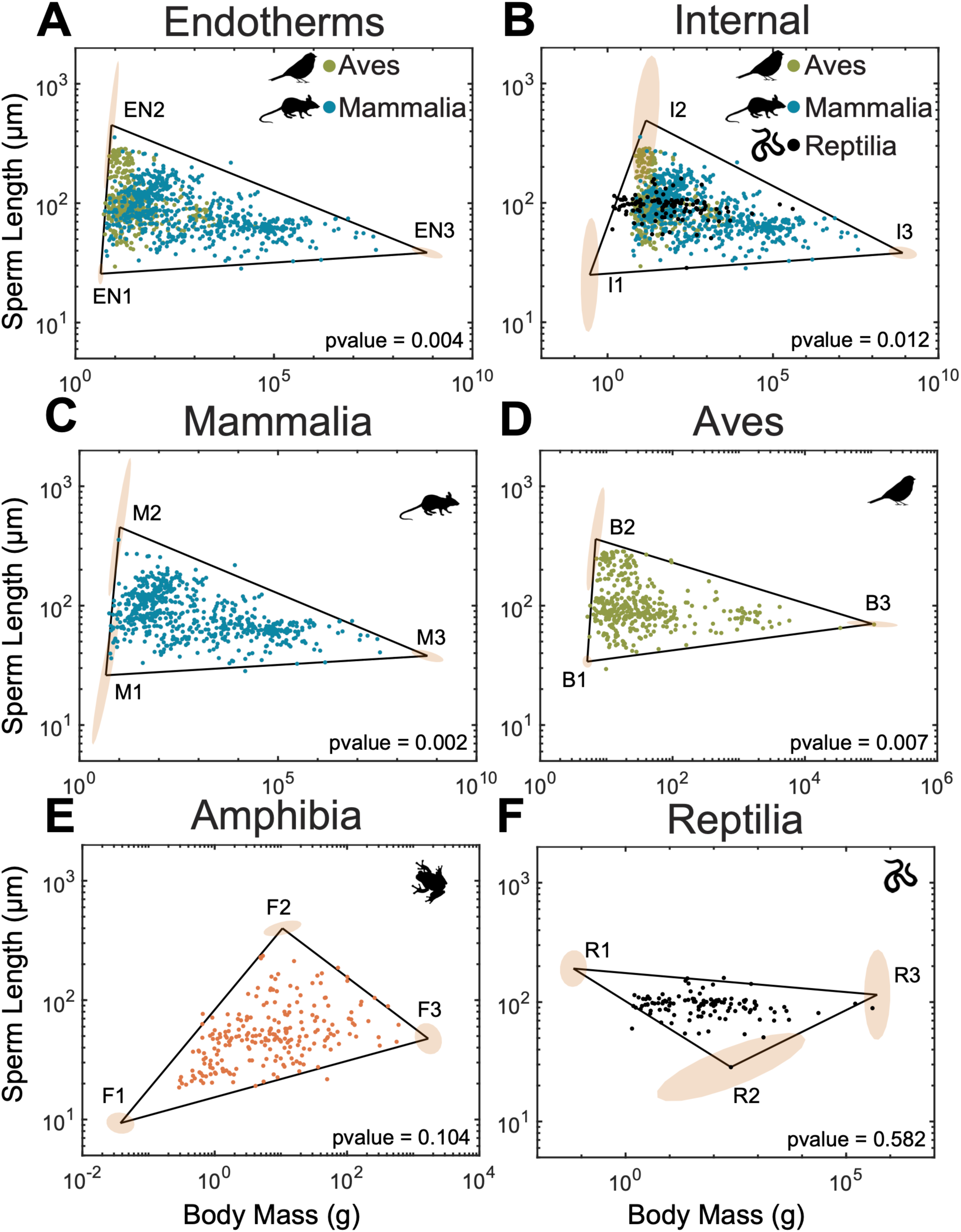
Robustness of the Pareto front within functional groups and class type. **A.** Endotherms, i.e., *Aves* and *Mammalia* (N=1042), and **B.** internal fertilizers, i.e., *Aves, Mammalia* and *Reptilia* (N=1157) show a triangular-shaped Pareto front (*p* = 0.004, *p* = 0.012 respectively). **C-D.** Mammalia (N=643), and *Aves* (N=399) show a triangular-shaped Pareto front (*p* = 0.002, *p* = 0.007 respectively). **E-F.** *Amphibia* (N=231) marginally show a triangular-shaped Pareto front (*p* = 0.104), while *Reptilia* (N=115) do not fall within a triangular Pareto front (*p* = 0.582).

**Figure 5.**
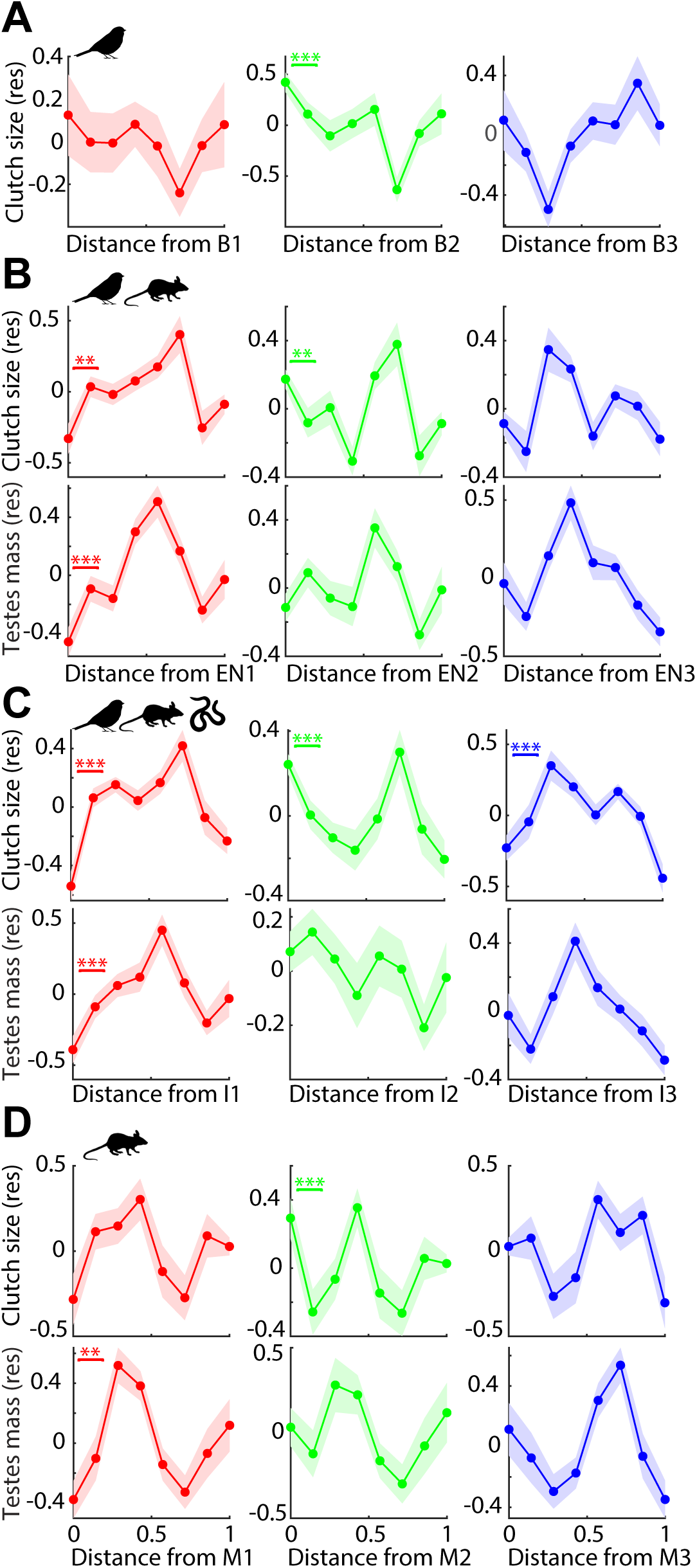
Feature density analysis for subgroups of tetrapods. We studied the feature density of clutch size and testes mass on **A.** *Aves*, **B.** Endotherms, **C.** Internal fertilizers and **D.** *Mammalia* by testing testes mass, clutch/litter size and genome size. We reported statistically significant features that enriched at least one vertex of the triangles in Fig. 4. Statistical comparisons were performed by running a Wilcoxon rank-sum test (\*\**p*<0.01, \*\*\**p*<0.001).

### Robustness of the Pareto front in relation to class type

To test the robustness of the Pareto front within each class, we grouped the species based on the four classes of tetrapods, namely mammals, reptiles, birds, and amphibians. We found that mammals and birds form significant sperm length-body mass triangular-shaped Pareto fronts (t-ratio test, *p* = 0.002, *p* = 0.007 respectively), while amphibians form a marginally not significant triangle (*p* = 0.104), and reptiles do not form a significant Pareto front (*p* = 0.582) (Fig. 4). Consistent with the case of endotherms and internal fertilizers, the vertex T2 in mammals and birds also enriched in clutch size (Fig. 5).

### Robustness of the Pareto front: control for phylogenetic bias

To discount the potentially confounding effect of phylogeny on the emergence of the triangular Pareto front, we used the flipping t-ratio approach^50^. This method entails randomly shuffling the values of either sperm length or body mass across phylogenetically close species while keeping unchanged the values of the other trait. This approach preserves both the phylogenetic constraints and the marginal distributions of body mass and sperm size.

We first controlled for phylogenetic dependencies at the class level. Since we had a different number of species for each class, we randomly selected 115 species, which corresponds to the maximum number of Reptilia we had in the dataset, for a total of 460 species. When shuffling the pairs of body mass and sperm size across species within each class, the triangular-shaped Pareto front remained significant (*p* = 0.021, Fig. 6), suggesting that our results are not affected by the phylogenetic dependencies at the class level.

**Figure 6.**
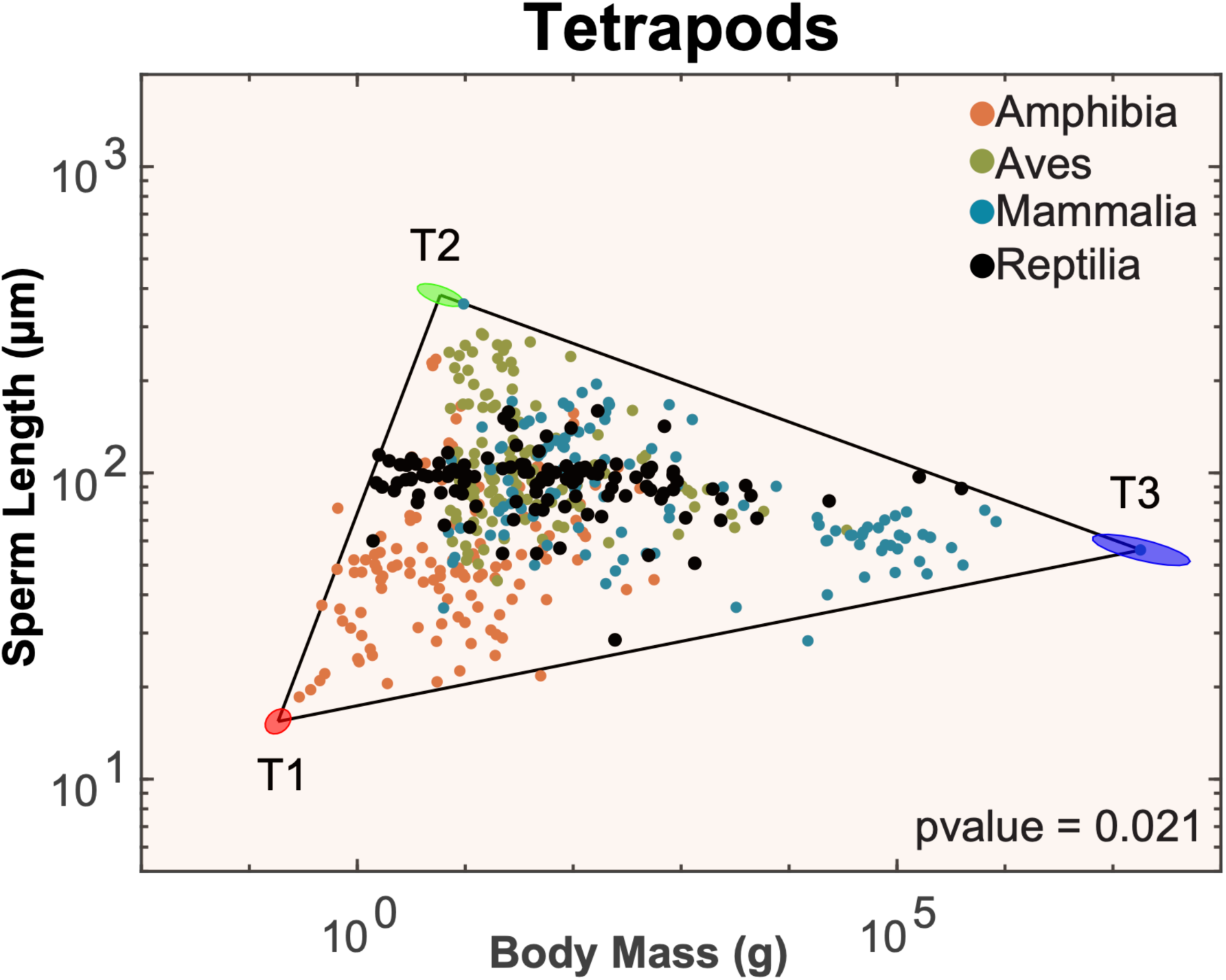
Robustness of the Pareto front: control for phylogenetic bias. We subsampled the initial dataset of 1388 species by picking up 115 species for each class in order to have the same number of species for each class, which resulted in a total of 460 species. We then used the flipping t-ratio approach to build the null distributions^50^. Data show a significant triangular Pareto front (t-ratio test, *p* = 0.021).

We next controlled phylogenetic dependencies at the order level. We considered only those orders that had at least 30 species, resulting in a total of 9 out of 35 orders. After shuffling the body mass and sperm size values across only those species within specific orders, we found that the Pareto front is still significant (*p* < 0.05). In conclusion, the Pareto front identified in tetrapods is robust to the taxonomic groups of mammals and birds, and against phylogenetic dependencies at both the class and order levels. Importantly, we also found that the Pareto front in tetrapods remains robust whilst down sampling to one-third of the data points from our tetrapod dataset (460/1388).

## Methods

### Data collection

For each species we collected a maximum of 5 trait values, namely body mass, sperm size, clutch size, testes mass, and genome size, that we defined as vectors *v*_*i*_ with *i* = 1, …, 5. We collected sperm size (µm) data from the most comprehensive published dataset to date on sperm size^47^ and from other sources e.g.^11,14,51^. Body mass, clutch size, testes mass as a proxy for sperm competition^9^, and genome size were collected from published and publicly available sources e.g.^52–56^. We included only a single source per species; when more than one value for a single species was present, we prioritized the most recent one. More details on our data collection procedure can be found in the Supplementary Materials and final sample sizes for each group and trait are reported in Table 1. Body masses and sperm sizes were log10-transformed before analysis. Testes mass and clutch/litter size are correlated with body mass, though in different directions among classes, and we thus obtained within-class residuals from log-log linear regressions between testes and body mass and between clutch size and body mass.

### The Pareto Optimality framework

Each species’ performance functions (*P*_*α*_(*v*)) depend on the specific combination of traits *v* for all tasks in trade-off (*α* = 1, …, *k* tasks). A basic assumption for the Pareto theory is the existence of a general fitness function, which is an increasing function of all the performance functions *F*(*P*_1_(*v*), …, *P_k_*(*v*)). Archetypes are the optimal phenotypes (*v*^*α*^) whose performance function *P*_*α*_(*v*^*α*^) is highest at a specific vertex *α*, which decreases as a function of distance from the vertex as *P*_*α*_(*v*) = *P̂*_*α*_((*v* − *v*^*α*^)^*T*^ *M*(*v* − *v*^*α*^)), where *α* = 1, …, *k* and M is a positive-definite matrix. As a result, no single species can excel at all tasks at the same time and the spatial coordinates of each species within the Pareto front provide information about the set of traits that will result in optimal Pareto solutions at any point along the fronts (Fig. 1).

### The triangularity test, density enrichment analysis and the position of the archetypes

We determined the statistical significance of the Pareto front’s triangularity with the t-ratio test (see Supp. Fig. 2). The t-ratio defines the ratio of the area of the best-fitting triangle that encloses the data to the area of the convex hull of the data. The closer the t-ratio is to one, the better the triangle captures the data distribution. We then compared the t-ratio of the original distribution with the ones obtained from 1000 randomized distributions. For each randomized distribution, we randomly shuffled the pairs of values for body mass and sperm size across phylogenetically close species^50^. This approach destroys the statistical correlations between the two traits while preserving the marginal distributions along single traits. The p-value was then defined as the fraction of times that the randomized distributions achieved a lower t-ratio than the original one. We considered as statistically significant p-values those that scored under 5% of times (*p* < 0.05).

According to the Pareto Optimality Theory, archetypal species near the vertices should manifest maximum values in a set of traits and decrease in their scores monotonically by increasing the Euclidean distance from the vertex. We followed the approach used in^46^ to analyze the feature density as a function of the Euclidean distance from the archetypes. For each feature we used the Wilcoxon rank sum test to test whether archetypes carried significantly higher values compared to the rest of the population^57^.

## Discussion

In this study, we postulated that the relationship between sperm length and body mass in tetrapods is shaped by trade-offs between multiple tasks. To test this hypothesis, we used the Pareto Task Inference (ParTI)^36^, a multi-objective optimization algorithm that can identify trade-offs among multiple tasks. We provide clear evidence that across tetrapods, sperm length variation in relation to body mass lies on a triangular-shaped distribution. According to Pareto Optimality theory, this suggests that different interacting evolutionary forces are simultaneously acting on the sperm length-body mass distribution. To infer which evolutionary factors may affect the sperm length-body mass phenotypic space, we explored the properties encoded by the species at the vertices of the Pareto front. We considered the three main factors that have been proposed to influence sperm evolution, namely the level of sperm competition, the number of eggs per clutch, and the size of the genome^2^. Our results reveal that these traits are either maximal or minimal at the vertices of the triangular Pareto front, overall demonstrating that body mass optimally shaped sperm size evolution in tetrapods mainly through its association with sperm competition and clutch size.

Our results unveil some important aspects of sperm size evolution in tetrapods. Firstly, our results show that sperm length variation is significantly associated with body mass, though not linearly. A negative association between sperm size and body mass has been postulated due to the predicted trade-off between sperm number and sperm size^58^, however, many studies failed to find a significant association between sperm size and body mass e.g.^14,29,59^. In contrast, we demonstrated that in tetrapods a nonlinear relationship exists. We found that sperm size is maximized at intermediate body size (T2), while is minimum in small animals (T1) and intermediate in large-bodied animals (T3). This result indicates that moving away from intermediate to small- or large-body sizes (from T2 to T1 or T3) limits the evolution of sperm size, indicating that sperm length-body mass relationship is mediated by trade-offs. Indeed, some of the key factors that have been postulated to impact sperm length evolution are linked – directly or indirectly – with body size and can generate opposite selective forces. Clutch size is expected to favour sperm size evolution^24,25^. In birds, for instance, clutch size is positively correlated with sperm size e.g.^11,23^, probably because larger sperm may be more efficient in entering into and persisting within the female storage organs, whose size is related to clutch size^60,61^. However, at the species level body size covaries with lifespan, which in turn is expected to be associated with a k-reproductive strategy and hence with small clutches, as observed for example in mammals, where very large species like ungulates, pinnipeds or cetaceans usually produce a single offspring^32^. In external fertilizers, in contrast, body size is positively correlated with clutch sizes, which in turn are expected to favour the production of numerous sperm, possibly at the expense of sperm size^51^. We found that sperm size is maximized in species with high sperm competition and, somewhat surprisingly, we found that sperm competition is maximized in species with intermediate body size (T2), whereas very small and very large tetrapods show a relatively low level of sperm competition (T1 and T3). On one hand, this result corroborates previous evidence which suggested that gametic investment is maximized in species with high level of sperm competition. However, the strength of this effect does not seem to increase as a function of body mass as postulated by the dilution hypothesis^32^. The fact that sperm size evolution is constrained at the extremes of the body mass range in tetrapods (moving from T2 to T3, body mass increases and sperm size decreases) may reflect the association between body size and ecological/social factors, such as population size, density and dispersal, which affect male and female reproductive strategies and hence levels of sperm competition^30–33^. Finally, we found that the size of the genome is maximized at the T3 vertex, which corresponds to animals with low-to-intermediate sperm sizes and large body masses, and it is minimized in species with long sperm and intermediate body masses (T2). This finding is particularly interesting, as it indicates that genome size does not represent a main driver in the evolution of long sperm and that genome size may be constrained in species with high sperm competition. This is because the cost of replicating a large genome may favour smaller genomes in species with high sperm competition which usually have a high rate of sperm production^62^ and/or the drag associated with a large sperm nucleus may reduce sperm swimming velocity, which provides an advantage in competitive fertilization success. Genome size, on the other side, may contribute to explaining why very large tetrapods have slightly longer sperm than very small ones (T3 *vs* T1), according to the hypothesis that larger genomes require longer sperm flagella to counteract the drag associated with a larger sperm head.

Finally, we revealed the association between body mass and sperm size across different taxonomic levels. We found that a triangular-shaped Pareto front is maintained by mammals, birds, endothermic species, and internal fertilizers, suggesting that similar evolutionary pressures characterize the evolution of sperm size in relation to body size within taxonomic/phylogenetic and functional subgroups within tetrapods. We highlight the power of the present statistical framework to detect complex, non-linear associations between traits. Unlike comparative analyses that typically control for body size variation^63^, the Pareto-optimality framework allows to investigate the evolution of sexual traits by explicitly including body size variation. We demonstrated that including body size variation can unveil previously unknown trade-offs among different factors, which can be inferred from the distribution of species in the trait space. However, we also acknowledge the limitations of our statistical approach. Firstly, the method requires larger sample sizes than comparative analyses to encompass the entire trait space determined by the evolutionary processes of interest. Further research is needed to confirm our results in subgroups, such as amphibians and reptiles, where Pareto fronts were not significant but species sampling was less numerous, thereby reducing the statistical power. Secondly, while this approach is useful for revealing previously unknown or untested trade-offs, it does not allow us to demonstrate causal relationships between the archetypes represented by traits enriching the vertices and the traits determining the Pareto front. For example, in our study, although testes mass and clutch size are both maximized in species with long sperm and intermediate small-to-intermediate body mass, our analysis does not allow us to disentangle whether large clutches have favoured the evolution of polyandry (and hence sperm competition and longer sperm), or if long sperm provide a fertilization advantage irrespective of the level of sperm competition, or if the two processes tend to covary in the same species because large clutches and polyandry are independently associated.

In conclusion, we present a comprehensive analysis of sperm size variation across tetrapods, based on the Pareto Optimality framework. All the hypotheses on the evolution on sperm length proposed so far assume that factors determining sperm length evolution are mediated, directly or indirectly, by body size. Congruently, our analysis demonstrated the existence of a triangular-shaped Pareto front in the space of body mass and sperm length, suggesting that these two traits reflect underlying evolutionary trade-offs and constraints influencing sperm length evolution. We showed that long sperm evolve in species with intense sperm competition, as expected^9^, and with large clutch size and we provided novel results that will promote future research. In particular, we demonstrated that the evolution of long sperm is constrained in very large and very small tetrapods, and this pattern is confirmed also across taxonomic and functional subgroups. The reasons why large clutch size and high levels of sperm competition are not observed in very small and very large tetrapods probably depend on different ecological and morphological constraints which need to be further investigated. We also demonstrated that long sperm do not occur in species with large genomes, suggesting genome size is constraining, rather than favouring the evolution of long sperm, contrary to predictions^27,28^.

## Acknowledgements

We thank Samuel Allan Kean for proofreading the article and Beniamino Tuliozi for comments on an early draft of the manuscript. We also would like to thank all the people that over the past decades collected the data that we used in this study and made it publicly available to the community. SC was supported by grants from University of Padova (BIRD-175144-2017) and by Valenzano lab core budget.

## Code and data availability

All data and the associated references are reported in the dataset uploaded in Figshare (temporary private link: https://figshare.com/s/61c3b2b6bf4e3f007894). We analysed the data using the ParTi code which is implemented as a Matlab software package available in Github (https://github.com/AlonLabWIS/ParTI).

## Author contributions

MBR, AM and AP conceived the study, SC and AP collected the data, LK performed the analyses and produced the results, LK and SC led the writing with contributions from all authors who approved the final version of the manuscript.

