## Supplementary materials for "Sperm length evolution in relation to body mass is shaped by multiple trade-offs in tetrapods"

### Supplementary methods: data collection

All data and the associated references are reported in the dataset uploaded in Figshare (temporary private link: https://figshare.com/s/61c3b2b6bf4e3f007894). During data collection, we prioritized data from recent sources that contained data for multiple species, and supplementing data for individual species when possible. To avoid conflicts among datasets, we followed a standardized protocol when collecting data. Firstly, we preferred sources that reported measures from n>1 individual and we thus excluded values measured from a single individual or from dead animals. We considered only mean values instead of maximal values. Then we preferred to not include values collected using data extraction software from images and values collected without a described and standardized methodology (e.g. "personal observations"). If multiple sources fit the above criteria, we prioritized more recent values. Where multiple data remained, we prioritized the dataset with the largest sample size. In a few cases we spotted clear errors in the values reported in the most recent study and we thus decided to report the value contained in the original reference. All the species names were uniformed to the most recent nomenclature or in the case of equivalent synonymous, the most used were chosen. For Amphibia, we limited our analysis to anurans species, thus excluding two orders: Urodela and Gymnophiona. While we did not find data relative to Gymnophiona species, we decided to exclude Urodela (salamanders) for two reasons: 1) on average their sperm are extremely long and significantly longer than the average sperm size of tetrapods[1] and 2) we did not found data on clutch size and testes mass for the salamanders in our dataset. Using salamander species for exploring sperm size-body mass morphospace would have extended the distribution of sperm size without the possibility to perform enrichment analysis on those phenotypes given the absence of clutch size and testes mass data. We also excluded an outlier for sperm size: an anuran species (*Discoglossus pictus*) in which males produce the longest vertebrate sperm measured (2.5 mm[1]). It is worth noting, however, that the exclusion of *Discoglossus pictus* and Urodela species did not significantly affect the shape of the morphospace given by the sperm size and body mass.

1 Birkhead, T. R., Hosken, D. J. & Pitnick, S. *Sperm Biology: An Evolutionary Perspective*. (Academic Press, 2009).

### Supplementary figures

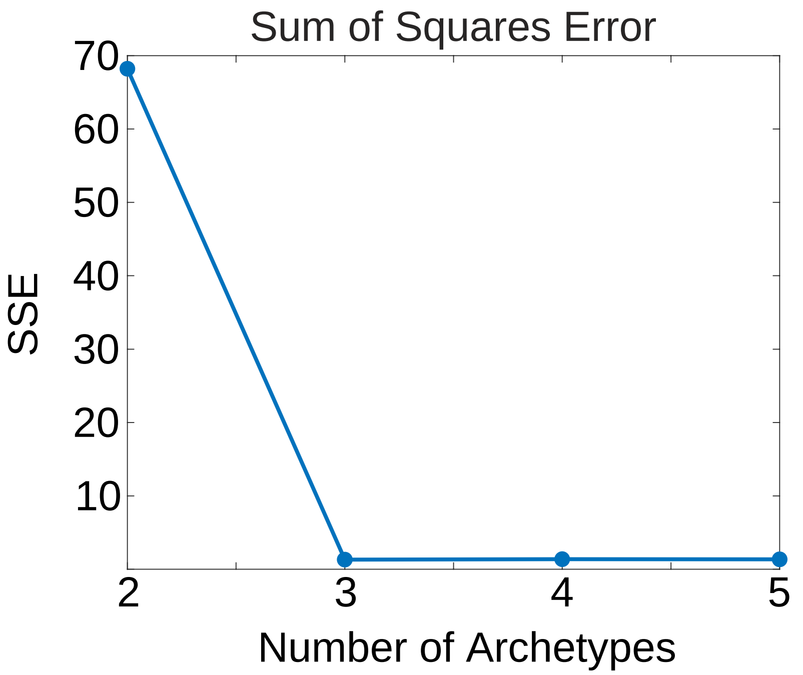


**Supplementary Figure 1. Sum of squares error as a function of the number of vertices.** Sum of squares error was computed with the PCHA algorithm. Three is the optimal number of vertices that minimizes the errors. Then, a plateau is reached.

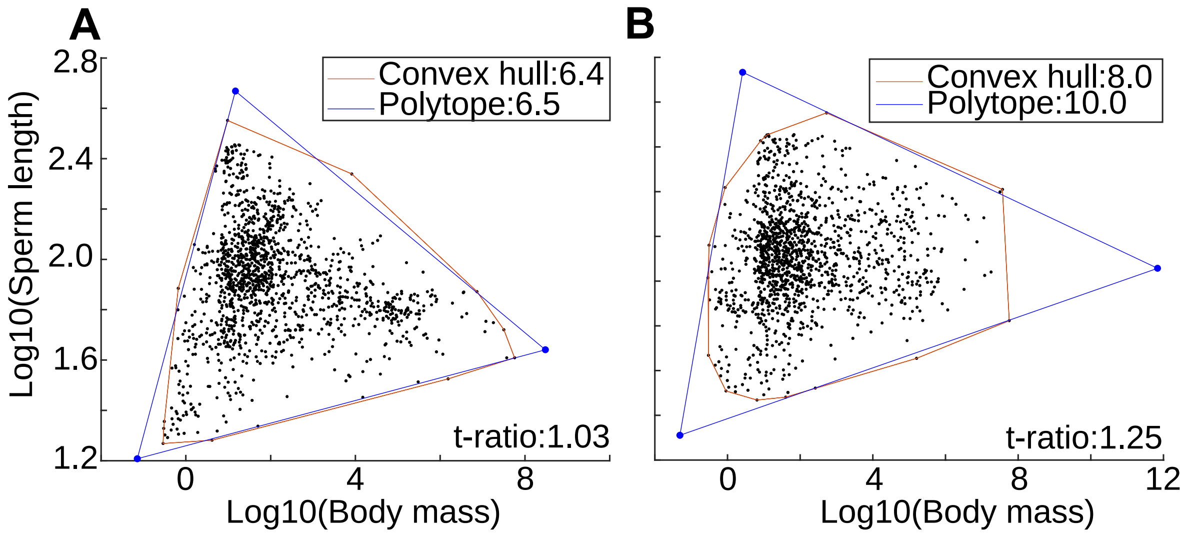


**Supplementary Figure 2. Test of triangularity to determine the significance of the triangular Pareto front. A.** Data points are shown as black dots. The red polygon corresponds to the best convex hull that encapsulates all the data points. The blue polygon represents the minimal triangle that encloses most data points but leaves some outliers outside. We computed it via the Sisal algorithm. The t-ratio is the ratio between the volumes of the triangular polygon and the convex hull. **B.** Randomized data points with the flipping t-ratio approach to correct for the phylogenetic influences.

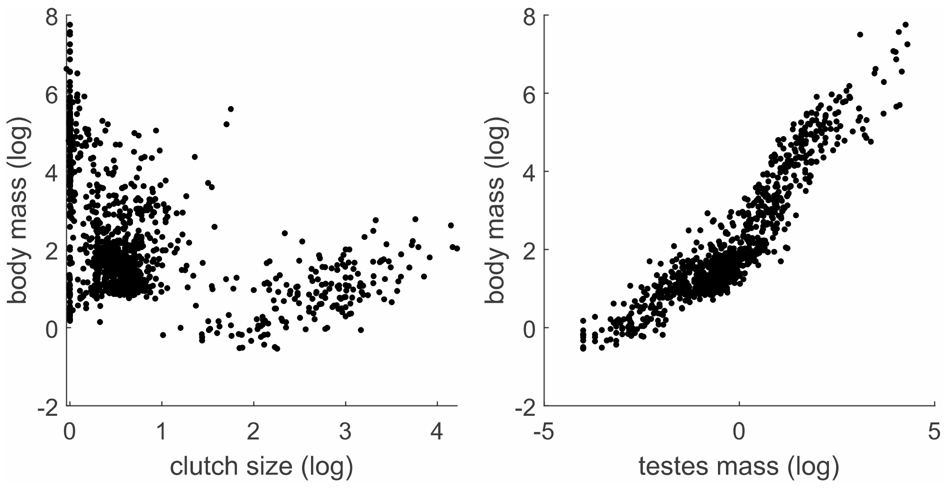


**Supplementary Figure 3. Relationship of body mass with clutch size and testes mass.** Tetrapods are plotted in the log-log space of body mass and clutch size in the left and in the log-log space of body mass and testes mass in the right.
